# High-throughput data-driven analysis of myofiber composition reveals muscle-specific disease and age-associated patterns

**DOI:** 10.1101/461608

**Authors:** Vered Raz, Yotam Raz, Davy van de Vijver, Davide Bindellini, Maaike van Putten, Erik Ben van den Akker

**Author notes:** Correspondence: Vered Raz, PhD, Leiden University Medical Center, Department of Human Genetics, Einthovenweg 20, 2333 ZC Leiden, Building 2, Room R3-17, The Netherlands. Shared senior authors (Maaike van Putten, Erik Ben van den Akker).

## Abstract

Contractile properties of myofibers are dictated by the abundance of myosin heavy chain (MyHC) isoforms. MyHC composition designates muscle function and its alterations could unravel differential muscle involvement in muscular dystrophies and aging. Current analyses are limited to visual assessments in which myofibers expressing multiple MyHC isoforms are prone to misclassifications. As a result, complex patterns and subtle alterations are unidentified. We developed a high-throughput data-driven myofiber analysis to quantitatively describe the variations in myofibers across the muscle. We investigated alterations in myofiber composition between genotypes, two muscles and two age groups. We show that this analysis facilitates the discovery of complex myofiber compositions and its dependency on age, muscle type and genetic conditions.

#### Nonstandard abbreviations

LGMD: Limb girdle muscular dystrophy
MFI: mean fluorescence intensity
MyHC: Myosin heavy chain
SGCA-null: α-sarcoglycan-null
SGCD-null: δ-sarcoglycan-null

## Introduction

Contraction properties of skeletal muscles are highly dynamic and may change in response to physical exercise, hormones, aging, or muscle diseases. Muscles are built of myofibers that differ by the expression of Myosin Heavy Chain (MyHC) proteins. In mouse muscles, the MyHC-1, MyHC-2a and MyHC-2b are the most abundant and are used to determine the contractile characteristics of myofibers (1, 2). Alternations in myofiber composition have been reported in several muscular dystrophies and myopathies, including Duchenne muscular dystrophy (DMD)(3-6), Oculopharyngeal muscular dystrophy (OPMD) (7), limb-girdle muscular dystrophy (LGMD) and other muscular dystrophies (8). Classically, myofiber characteristics are determined based on a visual assessment of MyHC expression, and are often categorized on basis of the expression of a single and most dominant isoform only (7, 9-11). In addition, visual assessments are subjective the number of myofibers that can be scored is limited. Moreover, myofiber classification that is based on binning of MyHC expression levels poorly capture the fully natural variations of myofiber heterogeneity. For instance, hybrids, expressing more than one MyHC isoform, are therefore often missing or misclassified (12). As a result, the variations that could be most relevant to muscle function and disease conditions are poorly captured.

Here we applied techniques from the image analysis domain to overcome limitations of myofiber typing by visual assessments. By automating routines for data acquisition, we drastically increase the number of myofibers analyses per sample to > 800 individual myofibers per tissue section. We next exploit the resulting richness of data points by training and applying models for an unbiased data-driven description of myofiber variation. We show that our high-throughput data-driven analysis facilitates the discovery of complex myofiber compositions and their dependency on age, muscle and genetic composition.

## Materials and Methods

### Animals and tissue preparation

Alpha-sarcoglycan deficient mice (B6.129S6-Sgca^tm2Kcam^/J (SGCA-null)) (13) were kindly provided by Queensta Millet, University College London and delta-sarcoglycan deficient mice (B6.129-Sgcd^tm1Mcn^/J (SGCD-null)) (14) were obtained from the Jackson Laboratory (Bar Harbor, ME, USA). C57BL/6J mice were used as wild types. Mice were bred in the Experimental Animal Facility of the Leiden University Medical Center where they were kept in individually ventilated cages with 12 hour light/dark cycles at 20.5°C and had *ad libitum* access to standard RM3 chow (SDS, Essex, UK) and water. Five C57BL/6J, SGCA-null and SGCD-null males aged 8 or 34 weeks were euthanized by cervical dislocation and the diaphragm and gastrocnemius muscles were dissected and snap frozen in 2-methylbutane (Sigma Aldrich, The Netherlands) cooled in liquid nitrogen. Samples were stored in −80°C. The experiment was approved by and performed following the guidelines of the Animal Experiment Committee (DEC 13211) of the Leiden University Medical Center.

### Antibody preparation

Hybridoma lines BA-D5, SC-71 and BF-F3 (Developmental Studies Hybridoma Bank (DSHB, USA) were kindly donated by Dr. Jennifer Morgan (UCL London). Antibodies produced by BA-D5, SC-71 and BF-F3 clones detect MyHC-1, −2a or −2b, respectively (15). Cells were cultured, and antibodies were purified as described in (16) with the exception that the sodium azide preservative was excluded from the supernatants. Supernatant antibodies were concentrated over IgG (BA-D5 and SC-71) or IgM (BF-F3) HiTrap columns (GE Healthcare Life Sciences, The Netherlands). Purified antibodies were conjugated to Alexa-350 (BA-D5), 594 (SC-71) or Alexa-488 (BF-F3) using the corresponding Protein Labelling Kits (ThermoFisher Scientific, Pittsburgh, PA, USA).

### Histology and Imaging

Muscle cryosections, 8 μm, were made with a Shandon cryotome (ThermoFisher Scientific) on Superfrost Plus slides (ThermoFisher Scientific). Haematoxylin and Eosin (H&E) staining was performed according to the manufacturer’s instruction and slides were mounted with Pertex (Histolab, Västra Frölunda, Sweden). Images were taken with a 10x objective with a BZ-X700 microscope (Keyence, Japan). Representative images are shown in Fig. 1A.

**Figure 1.**
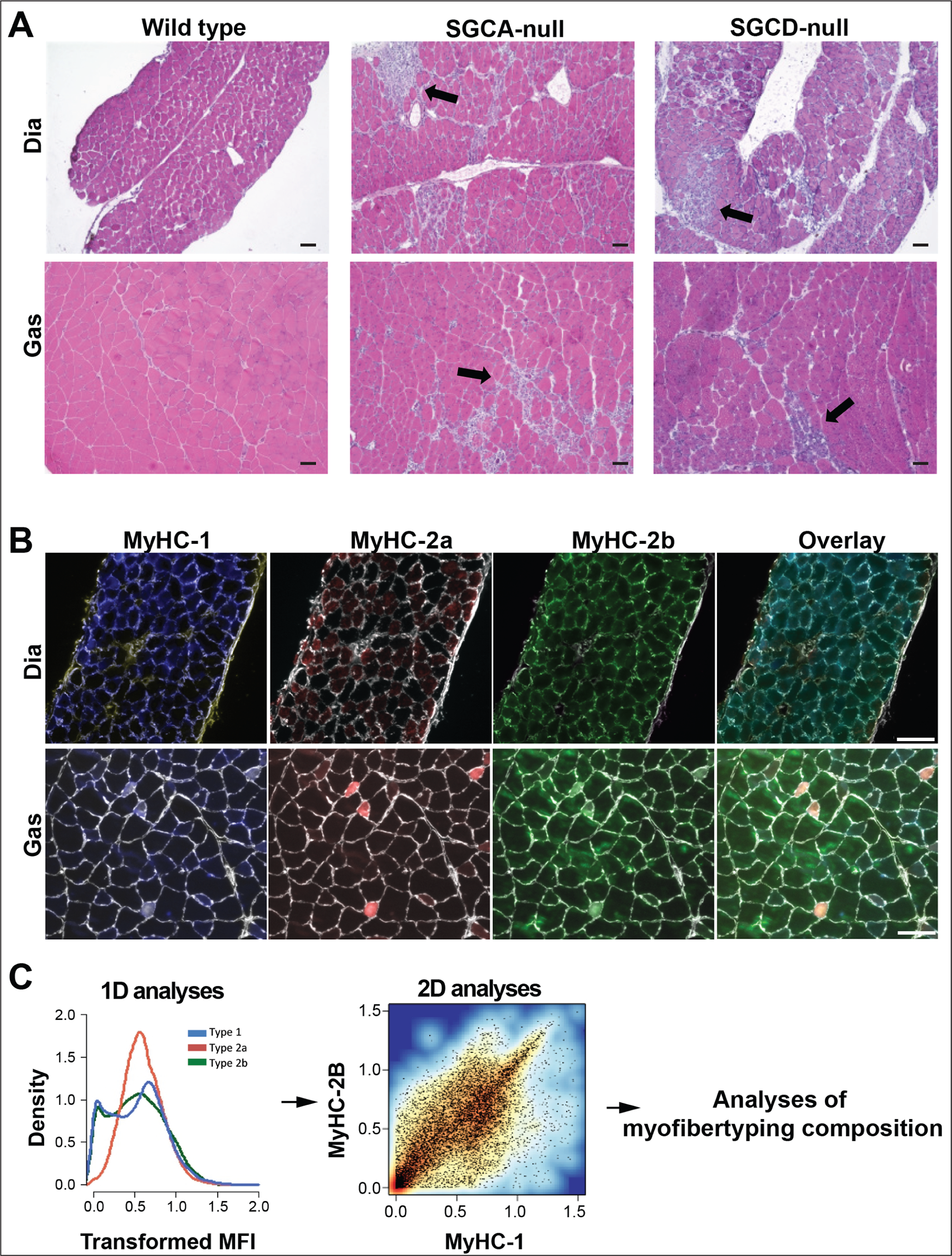
Histopathology and overview of the analysis pipeline. **A.** Representative images of muscle cross-sections of the diaphragm (Dia) and gastrocnemius (Gas) of 8-week-old wild type, SGCA-null and SGCD-null males. Images show H&E staining and arrows indicate regions with degeneration, regeneration, inflammation and fibrosis. Scale bar is 75 µm. **B.** A representative image of MyHC isoforms staining in diaphragm and gastrocnemius wild type muscles. Color code for MyHC isoforms is: MyHC-1 in blue, 2a in red and 2b in green. Laminin staining is depicted in white. Laminin was used for myofiber segmentation and for each myofiber MFI from each MyHC isoform was measured. Scale bar is 75 µm. **C.** Myofiber-based analyses of MyHC MFI include 1D analysis for individual MyHC isoforms, 2D analyses for two MyHC isoforms and quantitative analyses of myofiber distribution. Analyses were carried out on the pooled data from N=5 mice per genotype.

For immunofluorescence, slides were air dried for 30 minutes and circles were drawn around the tissue with a DakoCytomation pen (DakoCytomation, Denmark) followed by washing in PBS/0.05% Tween (PBST). Blocking was performed with 5% milk in PBST for 30 minutes followed by PBST washing. Sections were incubated with primary antibody (laminin 1:100, Abcam ab11575, USA) for three hours and after PBST washing sections were incubated with an anti-rabbit secondary antibody conjugated to Alexa 647 (1:1000; Invitrogen) for 90 minutes, following PBST washing MyHC isoforms immunofluorescence was carried out with fluorophore-conjugated monoclonal antibodies to: type-1_alexa-350 (1:75), type-2a_alexa-594 (1:800) and type-2b_alexa-488 (1:600). An overnight incubation was followed by extensive washing with PBST and mounting with ProLong Gold (Invitrogen). Images were made with a DM5500 fluorescent microscope (Leica, Wetzlar, Germany) using LAS AF (Leica) software V2.3.6. Per mouse muscle, five to nine images, representing the whole cross-sectional muscle area, were taken with a 10x objective. Tissue overlap was avoided. Representative images are shown in Fig. 1B.

### Image quantification

Single-myofiber quantification of MyHC isoforms mean fluorescence intensity (MFI) was performed employing a macro in STACKS V1.0, as previously described (17). In brief, single myofibers were segmented using a laminin staining. When laminin was non-continuous a line was extrapolated as ultimate erosion by shrinking the areas into a closed object and then expanding it to its original size. Regions consisting of incomplete fibers, folded tissue, gaps, freezing damage, blood vessels and connective tissue, were excluded from the analysis. Subsequently, the corresponding MFI values for each of the three MyHC-isoforms were collected from single myofibers. The background was subtracted, per image, by the macro. Prior to analysis MFI values were corrected for the fluorophore emission efficiency that is specific for the microscope (the specific values are: 430 nm = 0.735; 488 nm = 0.067; 596 nm = 0.172).

### Analysis of myofiber composition

Myofiber composition was investigated using 1D and 2D density plots. For this purpose, MFIs of each MyHC-isoform were pooled across fibers from all mice per condition: muscle type, age and genotype and then scaled per sample (without centering). 1D and 2D density plots were created using the misc3d package in R. The best of the regression line was calculated using a linear regression model. Gating was determined from the 2D density plot, the number of myofiber within a gate was counted and the proportion was calculated from the pooled number of myofiber per condition. Subpopulation analysis of myofibers was performed with a density-based clustering using the mean shift algorithm, implemented in the LPCM package in R. Given a bandwidth H, indicating a measure of proximity between data points, mean shift algorithm identifies local maxima in the data by iteratively shifting proximate data points towards areas of higher density. At each iteration, points are shifted towards the steepest density gradient and densities are recomputed, until all points have merged into a local maximum. Hence, this procedure will simultaneously yield the local maxima of density and the points that contributed to it, i.e. a clustering of data points. Mean shift was run using an increasing bandwidth, (H ranges 0 to 0.20) and cluster sizes were recorded. To assess the homogeneity in 2D data, we computed the percentage of myofibers assigned to the largest cluster (%LC). %LC near zero represents no major cluster and %LC=100 indicates the presence of one dominant cluster. To compare between different biological conditions the number of myofibers subjugated to the repetitive clustering were down-sampled to the lowest number of fibers (N=5173). Trend lines were fitted using the top 5% densest areas. For this purpose, data was assigned to a raster of 100 x 100 equally sized bins. The positions of the 5% bins with the most data points (e.g. 50) were then used for fitting a trend line with its intercept forced through the origin.

## Results

To assess alterations in myofiber composition we studied two muscles, the diaphragm and gastrocnemius, and three genotypes C57BL/6J wild type and two mouse models for LGMD type-2D and −2F, having mutations in α-sarcoglycan (SGCA-null) and δ-sarcoglycan (SGCD-null), respectively (13, 14). We first investigated muscles from 8-week-old mice, as at that age dramatic differences in muscle physiology were found in both mouse models as compared with wild type mice (13, 14, 18). We confirmed muscle pathology in SGCA-null and SGCD-null diaphragm and gastrocnemius, using H&E staining, and observed regions with degeneration, regeneration, inflammation and fibrosis (Fig. 1A). Both muscles were selected for this study since they express MyHC isoforms at different levels: the gastrocnemius predominantly expresses MyHC-2b and has lower expression levels of MyHC-2a and 1, whereas the diaphragm has higher levels of MyHC-2a and MyHC-1 (19, 20). Myofiber staining was carried out on muscle cross-sections with a mixture of antibodies to MyHC-1, −2a and −2b isoforms and anti-laminin. Laminin was used to segment the individual myofibers. Representative images are shown in Fig. 1B. Using a semi-automated procedure for myofiber-based image quantification, the mean fluorescence intensity (MFI) per individual myofiber was recorded across all sections. Per mouse, data was pooled from five mice, the number of myofibers per mouse and muscles are summarized in Fig. S1. To overcome the differences in the number of myofibers between individual mice (Fig. S1) we pooled myofibers per condition. This resulted in an enhanced power in the analyses. Per analysis >5000 myofibers were included (Fig. S1). We first analyzed the data using 1D distribution plots and subsequently clusters were identified using 2D analyses (Fig. 1C).

### 1D density plots reveal muscle and genotype specific alterations in myofiber composition

1D density plot in the diaphragm of 8-weeks-old wild type mice. MyHC-1 and MyHC-2b MFI showed a bimodal distribution (Fig. 2A), indicating the presence of two major myofiber populations: one with high expression levels and one with low expression levels of these MyHC isoforms. Despite the expected variations between mice, bimodal distributions were consistent across mice (Fig. S1). In contrast, the distribution of MyHC-2a was unimodal, indicating a single population, which did not differ between mice (Fig. 2A and Fig S1). In the gastrocnemius, MyHC-1 and MyHC-2b MFI distribution was predominantly unimodal, despite variations between mice (Fig. 2A, Fig. S1). The distribution of MyHC-1 and MyHC-2b was shifted to lower MFI in both LGMD models in both muscles (Fig. 2A). However, in the diaphragm of SGCD-null the shift to low MFI was more pronounced compared with SGCA-null (Fig. 2A). In the gastrocnemius the change in MyHC-2b was more pronounced compared with MyHC-1 (Fig. 2A). Together, this suggests that myofiber composition is differentially affected between diaphragm and gastrocnemius muscles in both LGMD models. Density plots of MyHC-2a were unchanged between muscles and between genotypes (Fig. 2A), therefore, MyHC-2a was excluded from the 2D analysis.

**Figure 2.**
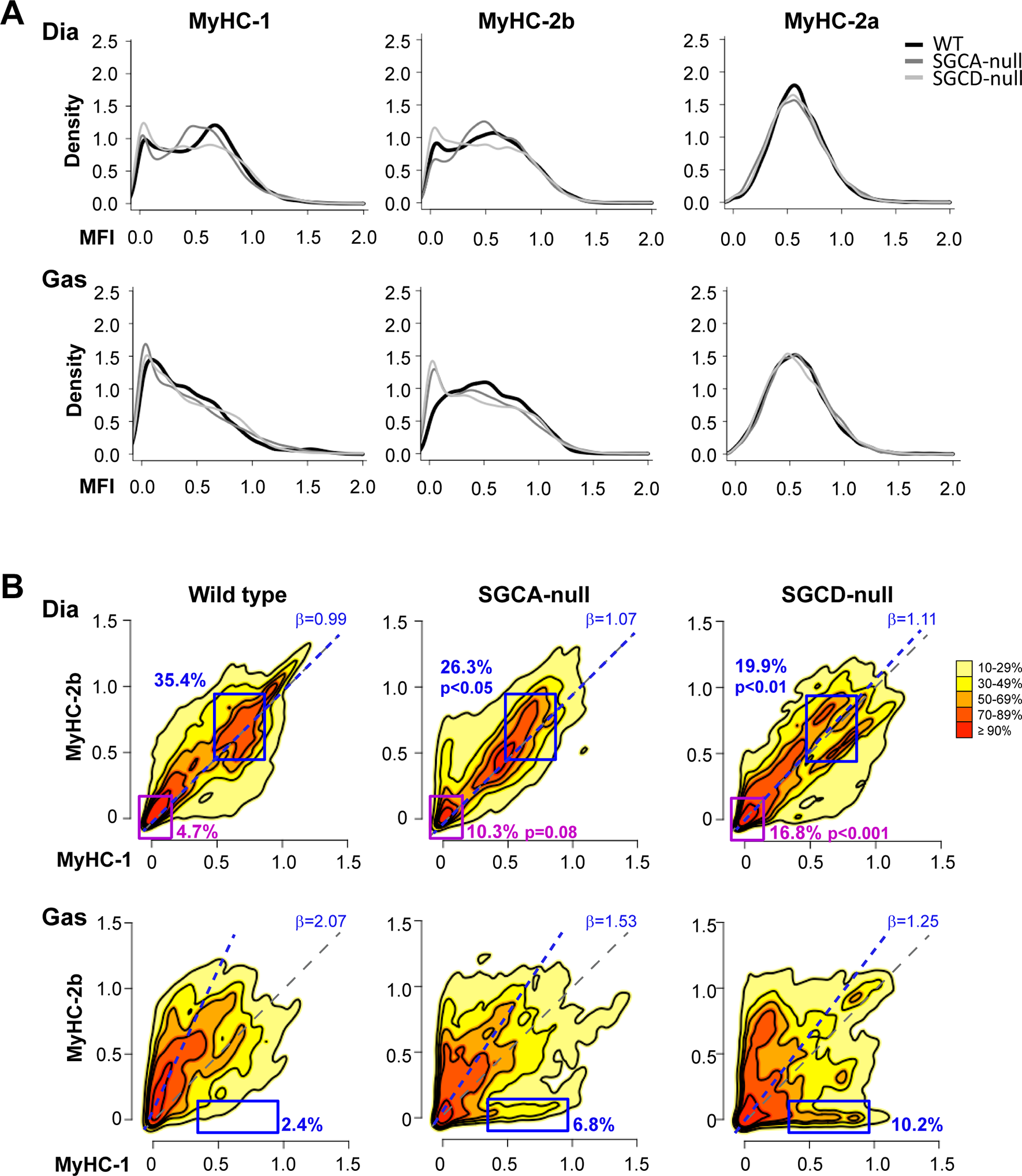
MyHC isoform distributions in diaphragm and gastrocnemius in wild type and LGMD mouse models. 1D and 2D plots were generated from the pooled myofiber data generated from 8-week-old mice per muscle per genotype. **A.** 1D density plots of MFI from the pooled data are per MyHC isoform, wild type is in black and LGMD mouse models are in grey. **B.** 2D density landscape plots of the pooled data. Colors indicate data density; the highest density (>90%) is depicted with dark red, and the least dense (10%) with light yellow. The purple and blue rectangles gate the high density areas in wild type diaphragm (Dia), and the same coordinates are projected in the 2D plots of SGCA-null and SGCD-null mice. The percentage of myofibers within the purple or blue areas is depicted with a corresponding color. P-value is calculated by the student’s t-test from wild type. In the gastrocnemius (Gas), the blue area gates a novel subpopulation in the SGCD-null and the same coordinates are projected in SGCA-null or wildtype 2D plots. The diagonal is depicted with a gray dashed line, the regression is depicted with a blue dashed line, and the beta is indicated in the upper right corner.

### 2D analyses reveal myofiber populations that differ between LGMD models

To further investigate the differences in myofiber composition between wild type and LGMD models we examined MFIs of MyHC-1 and MyHC-2b using their joint density 2D distribution. To achieve a power in 2D analyses, analyses were performed on the pooled data from N=5 mice and all myofibers were included. Using a 2D landscape plot, two density centers of myofibers were found in the diaphragm of wild type mice (Fig. 2B). The cluster with low MyHC-1/MyHC-2b MFIs included 4.7% of total myofibers (the purple gated area, Fig. 2B, Dia), while the group with high MFIs for both isoforms included 35.4% of total myofibers (the blue gated area, Fig. 2B, Dia). The two density centers were aligned with the diagonal (beta of the linear regression=0.99, blue dashed line Fig. 2B, Dia). This indicates a linear relation between MyHC-1 and MyHC-2b MFIs. The joint density distribution in both LGMD models differed from the wild type. Most notably, the subpopulation with high MyHC-1/MyHC-2b MFI values was shifted to lower values in the SGCA-null mice, while in the SGCD-null model this cluster became disperse (Fig. 2B, Dia). For a quantitative assessment of alterations in myofiber composition in the LGMD mouse models, we determined the percentage of myofibers within the purple or blue gated areas. In the SGCA-null model, the percentage of myofibers that are associated with both gated areas was only significantly decreased for the blue gated area (26.3%), while the increase in the purple area was non-significant (10.3%)(Fig. 2B). Notably, in the SGCD-null model the percentage of myofibers within the purple area significantly increased (16.8%), but the percentage of myofibers within the blue area significantly decreased (19.9%) (Fig. 2B, Dia, SGCD-null). This indicates that myofiber composition is highly changed in the SGCD-null model but only moderately changed in the SGCA-null model. The beta in SGCA-null and SGCD-null models was unchanged (beta=1.01; 1.11, respectively). This suggests that the relation between MyHC-1 and MyHC-2b is unchanged.

2D plots of the gastrocnemius in SGCA-null and SGCD-null mice revealed a novel group with high MyHC-1 and low MyHC-2b MFI values, which was not observed in wild type mice (Fig. 2B, Gas, gated in a box). The percentage of myofibers in this group was higher in SGCD-null compared with SGCA-null mice (10.2% and 6.8%, respectively, Fig. 2B, Gas). In addition, the relation between MyHC-1 and MyHC-2b MFI also altered in the LGMD mouse models, as compared to wild type mice. In the wild type mice beta of the regression line was 2.07, but in SGCA-null and SGCD-null models betas were 1.53 and 1.25, respectively. This indicates that in wild type MyHC-2b is expressed at higher levels compared with MyHC-1, but in the LGMD models MyHC-2b reduced whereas MyHC-1 increased. The reduced beta is consistent with the presence of the novel myofiber group with high MyHC-1 and low MyHC-2b MFI values in the LGMD models. Together, the 2D analyses in gastrocnemius suggest a trend in alterations of myofiber composition between these LGMD models with more severe changes in the SGCD-null model compared with the SGCA-null model. In the diaphragm alterations in myofiber composition are predominantly along the diagonal, whereas in the gastrocnemius changes occur along the axes. This suggests that different regulatory pathways affect myofiber compositions in the two muscles.

### Myofiber sub-populations change with age in wild type muscles

We next assessed whether age affects myofiber composition. We generated myofiber MFIs datasets in both muscles from 34-weeks old wild type mice using the same procedure as for the 8-weeks old mice. 1D MFI distribution age-associated changes were observed for MyHC-1 and MyHC-2b in the diaphragm of wild type mice, but were less profound in the gastrocnemius (Fig S3). The density distribution of MyHC-2a was predominantly unchanged with age in both muscles (Fig. S3). Therefore, 2D analyses were carried out only for MyHC-1 and MyHC-2b, similar to the 8-weeks old mice. The joint density distribution of the diaphragm revealed that the two major myofiber clusters were merged into one cluster with age (Fig. 3A, Dia). The single cluster in the 34-weeks old mice had intermediate MFI values, compared with those in the 8-weeks old animals (Fig. 3A, Dia). To assess if the relation between MyHC-1 and MyHC-2b was affected by age we calculated the beta of the regression line, trained on data of the top 5% densest areas. In the diaphragm the regression line overlapped with the diagonal, this indicates a symmetric relation between MFI values of MyHC-1 and MyHC-2b. In the gastrocnemius, the regression line showed a higher angle compared with the diagonal. This indicates lower expression of MyHC-1 compared with MyHC-2b. This is in agreement with other studies showing higher MyHC-2b expression in the gastrocnemius (19, 20). In both muscles the beta of the regression line was unchanged between the 8 and 34 weeks old mice (Fig. 3A, Dia: beta = 0.99 and 1.0, respectively, Gas: beta= 2.07 and 2.05, respectively). This suggests that the global relation between MyHC-1 and MyHC-2b MFI is unaffected by age in both muscles. Together, these results indicate that age-associated alterations in myofiber composition are prominent in the diaphragm and minor in the gastrocnemius.

**Figure 3.**
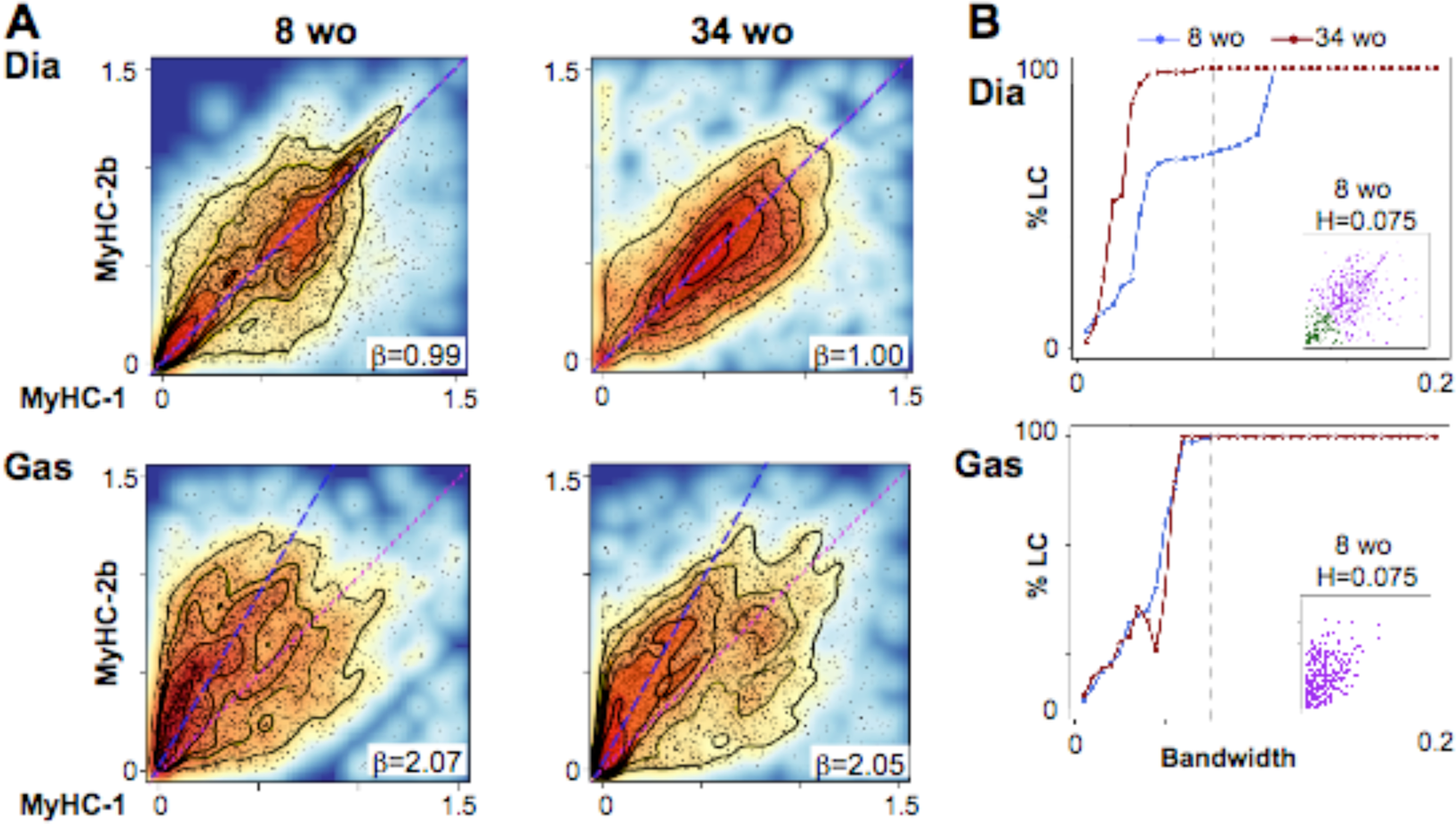
Age affects myofiber distribution in a muscle-specific manner. **A.** 2D distribution of MyHC-1 versus MyHC-2b MFI from the pooled data of the diaphragm (Dia), and the gastrocnemius (Gas) of wild type mice at 8 or 34 weeks old (wo). Each scatter plot is overlay with a corresponding topography plot. The >70% density regions were used to calculate a correlation between MyHC-1 and MyHC-2b. The diagonal is depicted with a purple dashed line and the regression is depicted with a blue dashed line. The beta is denoted in right bottom corner. **B.** Myofiber-type clusters. Plots show the percentage of myofibers that are assigned to the largest cluster (y-axis) across a fixed bandwidth range (x-axis) in diaphragm (Dia), or gastrocnemius (Gas) at 8 (blue) or 34 (red) weeks of age. Each point represents the result of a density-based clustering for a defined bandwidth and the line connecting the dots shows the trend. A bandwidth H=0.075 is depicted with a dotted line. The scatter plot shows clustering assignment for a bandwidth H=0.075 in the 8 weeks-old (wo) wild type mice.

### Myofiber heterogeneity is altered by age and genetic conditions

To quantitatively assess the age-associated alterations in myofiber subpopulations we performed mean shift clustering within a range of bandwidths (smoothness of estimated densities) and compared clustering results between 8-weeks and 34-weeks old wild type mice (Fig. S4). The percentage of fibers that were assigned to the largest cluster (%LC) was used as readout for heterogeneity in myofibers. In the diaphragm of the older mice 100% of the fibers were stably assigned to the largest cluster already from 0.045 bandwidth (Fig 3B, Dia), indicating that more than one subpopulation was robustly detected. In contrast, in the younger mice, a single stable cluster was found only from 1.2 bandwidth. In the range between 0.045-0.1 bandwidth only close to 65% of the fibers were assigned to the largest cluster whereas 35% were assigned to a second smaller cluster (Fig 3B, Dia). This demonstrates that two myofiber clusters, which are defined based on MyHC-1 and-2b MFI, are robustly identified in the young mice but only one cluster in the older mice. In the gastrocnemius, however, only one stable cluster was found, and it was indistinguishable between young and older mice (Fig. 3B, Gas). Together, this reveals that myofiber subpopulations differs between the two muscles in young mice, and the pattern changes with age in the diaphragm but not in the gastrocnemius.

We then also assessed myofiber heterogeneity in the LGMD models. Since in wild type mice the age-associated changes in clustering was more prominent in the diaphragm, we focused on that muscle. Two major clusters were found in the 8 week old SGCA-null mice, but in the older mice heterogeneity was reduced to only one cluster (Fig. S5). In the young SGCD-null mice only one dominant cluster was found, which remained a single cluster in the older mice (Fig. S5). Also in the older wild type mice only one cluster was found (Fig. S5). This suggests that myofiber heterogeneity in the diaphragm in the older LGMD models resembles to that in the wild type mice. Further this result suggests that alterations in the myofiber composition in dystrophic models reduce with age. This negative age-associated trend agrees with studies in the *mdx*mouse for Duchenne muscular dystrophy, where reduced muscle pathology was found in older mice (21, 22).

## Discussion

Skeletal muscles have the ability to shift myofiber composition in response to varied physiological and pathological conditions. So far, most myofiber analyses are limited to MFI binning, which are driven by the categories made by eye-assessment. This results in myofiber categories defining myofibers as negative or positive to a specific MyHC isoform (6, 7, 9, 12, 23-26). Hence, classical myofiber typing does not capture the full spectrum of myofiber variation, and most studies define myofibers type based on fast (expressing MyHC-2b) or slow (expressing MyHC-1/2a) twitch characteristics (9, 27). Here we define myofibers using a data-driven and hypothesis-free analysis, we show that the majority of myofibers express more than one MyHC isoform and the pattern of myofiber composition is distinguished between muscles. Application of semi-automatic image quantification can be combined with data-driven analysis of tens of thousands of myofibers (17, 28, 29). We considered the full range of MFI levels without binning of the data and show that myofiber composition is altered between muscles, wild type and LMGD models and with age. Most apparent changes were found in the diaphragm, suggesting that this muscle is more susceptible to age-associated alterations compared with the gastrocnemius. Moreover, we show that most prominently, myofiber heterogeneity is reduced with age. This suggests that a heterogenic myofiber composition characterizes young mice, and therefore could be beneficial. An age-associated differential muscle involvement was also reported in control human subjects and in wild type rats (11, 34), suggesting that the susceptibility for age-associated damage varies between muscles.

Exploring myofiber composition in mouse muscles, we found alterations between wild type and LGMD models. Enhanced alterations in myofiber composition were found in the SGCD-null mice compared with SGCA-null mice. This is consistent with a more severe muscle phenotype in SGCD-null mice compared with SGCA-null mice (13, 14, 18). In addition, we show differential muscle involvement between SGCA-null and SGCD-null mouse models, concomitant with differential muscle involvement in muscular dystrophies (30-33).

In this study analysis of myofiber composition is limited to MyHC-1, MyHC-2a and MyHC-2b. Here we focused on 2D analyses of MyHC-1 and 2b, since the distribution of MyHC-2a was not altered between genotypes or age. To capture additional myofibers subgroups more molecular features should be included. For example, future studies can include MyHC-2x, as it plays a role in muscle atrophy and aging (17), or neonatal and embryonic MyHC isoforms which are highly expressed in muscular dystrophy models (18, 35). In this study we did not address inter-individual variations, since N=5 has a limited power for 2D analyses. Pooling of myofiber MFI data allowed us to assess the robust changes in the mean but not in variation in 2D analyses. Our statistical assessment of specific changes in myofiber composition supports physiological studies showing that the SGCD-null model is more severe compared with the SGCA-null model. Specifically, changes in the diaphragm but not in the gastrocnemius were found significant. Future studies that are designed to study variations should statistically investigate the broad range of changes in myofiber composition.

Classification of myofiber composition within a muscle could be helpful to elucidate muscle cell biology prior to muscle degeneration or disuse. The underlaying mechanisms for myofiber composition remain obscure. Identification of clusters in myofiber composition is an initial step to understand the complexity of myofiber contractile features, from which experiments to unravel mechanisms regulating alterations of myofiber composition could be conducted.

## Acknowledgements

This work is funded by AFM grant 20251 for M. van Putten and AFM grant 21160 for V. Raz.

We thank M. Reinders (Dept of Biomedical Data Sciences, Leiden University Medical Centre, and Delft Bioinformatics Lab, Delft University, the Netherlands) and A. Aartsma-Rus (Dept of Human Genetics, Leiden University Medical Center, the Netherlands) for the helpful suggestions and comments on the analyses and the manuscript.

## Author contributions

V. Raz, M. van Putten and E.B. van den Akker designed the experiments

D. van de Vijver, D. Bindellini, M. van Putten performed the experiments

Y. Raz and E.B. van den Akker analysed data

V. Raz, M. van Putten and E.B. van den Akker wrote the paper

